# PT150 is a modulator of glucocorticoid and androgen receptors with antiviral activity against SARS-CoV-2

**DOI:** 10.1101/2020.07.12.199505

**Authors:** Neil D. Theise, Anthony R. Arment, Dimple Chakravarty, John M. H. Gregg, Ira M. Jacobson, Kie Hoon Jung, Sujit S. Nair, Ashutosh Tewari, Archie W. Thurston, John Van Drie, Jonna B. Westover

**Affiliations:** Department of Pathology, New York University-Grossman School of Medicine, New York NY USA; Palisades Therapeutics/Pop Test Oncology LLC, Cliffside Park NJ USA; Department of Biology, United States Air Force Academy, Colorado Springs CO USA; Department of Urology and the Tisch Cancer Institute, Icahn School of Medicine at Mount Sinai, New York NY USA; Department of Medicine, New York University-Grossman School of Medicine, New York NY USA; Department of Animal, Dairy, and Veterinary Sciences, Utah State University, Logan UT USA; ADME Solutions, Inc, San Diego CA USA; Van Drie Research, N. Andover MA USA

## Abstract

PT150 is a clinical stage molecule, taken orally, with a strong safety profile having completed Phase 1 and Phase 2 clinical trials for its original use as an anti-depressant. It has an active IND for COVID-19. Antiviral activities have been found for PT150 and other members of its class in a variety of virus families; thus, it was now tested against SARS-CoV-2 in human bronchial epithelial lining cells and showed effective 90% inhibitory antiviral concentration (EC_90_) of 5.55 μM. PT150 is a member of an extended platform of novel glucocorticoid receptor (GR) and androgen receptor (AR) binding molecules. In vivo, their predominant net effect is one of systemic glucocorticoid antagonism, but they also show direct downregulation of AR and minor GR agonism at the cellular level. We hypothesize that anti-SARS-CoV-2 activity depends in part on this AR downregulation through diminished TMPRSS2 expression and modulation of ACE2 activity. Given that hypercortisolemia is now suggested to be a significant co-factor for COVID-19 progression, we also postulate an additive role for its potent immunomodulatory effects through systemic antagonism of cortisol.

## Introduction

PT150, developed initially for treatment of depression (completing Phase 2 clinical trials), is both a down-regulator of androgen receptor (AR) expression (1) and a potent glucocorticoid receptor (GR) antagonist (2, 3). It is one of a suite of related compounds with similar hormonal receptor signaling functionalities which have demonstrated antiviral activity in a number of viruses across many virus families (unpublished data). Some effects are mediated by binding to viral genomic glucocorticoid response elements (GRE’s; e.g. in HIV, Ebola, HBV, HCV) (4). Many viruses without GRE’s have also been found to be potently inhibited by members of the platform. Two compounds, PT150 and PT156 were found to have some degree of in vitro antiviral activity against MERS virus. PT150, a clinical stage oral medication with a strong safety profile (5, 6) and an active IND for COVID-19, is now shown to have potency against SARS-CoV-2.

Proposed mechanisms of action for the potential suppression of these viruses by these compounds and, in particular, for treatment of the clinical syndrome of COVID-19 include direct antiviral effects at the virus/cell level and systemic effects arising from the inflammatory modulating activities related to their potent antagonism of cortisol in humans. The direct antiviral effects may relate to AR suppression through subsequent down regulation of transmembrane serine protease 2 (TMPRSS2), an androgen-dependent membrane receptor that primes the SARS-CoV-2 spike protein, and ACE2, the major cell receptor for viral entry (7–10). Moreover, systemic elevation of circulating cortisol (11) and virus induced neuro-inflammation (12) are now considered possibly significant co-factors in COVID-19 disease severity and progression. The combination of antiviral and immune modulatory effects may provide significant therapeutic value for COVID-19 patients.

## Methods

### MERS

#### Compounds

PT150 and PT156 were provided as a solid and stored at 4 °C upon arrival. Just before the assay, the test compounds were dissolved in DMSO to 20 mg/mL. Eight half-log, serial dilutions in culture medium Minimum Essential Medium supplemented with 5% FBS and 50 μg/mL gentamicin with a highest test compound concentration of 100 μg/mL.

#### Cell culture

96-well plates were seeded with MA-104 cells purchased from ATCC, MA-104 Clone 1 (ATCC^®^ CRL-2378.1) and incubated overnight. Seeding was performed at a cell concentration that yielded 80-100% confluent monolayers in each well after overnight incubation. The test media used was the culture medium with 5% FBS and gentamicin.

#### Virus

MERS strain EMC was passaged twice in MA-104 cells to produce the stock. Virus was diluted in the test medium with 5% FBS and gentamicin. The multiplicity of infection (MOI) was 0.001 CCID_50_ per cell.

#### Experimental design

100 μL of each concentration were added to 5 test wells on the 96-well plate. Three wells of each dilution were treated with the test virus in test medium. Test medium with no virus was added to 2 wells as uninfected toxicity controls. Six wells were untreated virus controls. The protease inhibitor M128533 (E128533; Epicept Corp, San Diego CA), a known compound with activity against MERS, was assayed in parallel (13).

The plates were incubated at 37 °C + 5% CO_2_ until cytopathic effect (CPE) was apparent in the virus control wells. After CPE was observed microscopically, wells were stained with 0.011% neutral red dye for approximately 2 hours. Dye was then siphoned off and 200 μL 50:50 Sorensen citrate buffer/ethanol for was added for >30 min., agitated, and read on a spectrophotometer at 540 nm. Optical density was converted to a percent normalizing to the cell control wells. EC_50_ and CC_50_ were calculated by regression analysis.

### SARS-CoV-2

#### Compounds

The test compound was provided as a solid and stored at 4 °C upon arrival. Just before the assay, the compound was dissolved in 100% DMSO at a concentration of 20 mg/mL and further diluted to the test dilutions in the MatTek culture medium (AIR-100-MM).

#### Cell Culture

Differentiated normal human bronchial epithelial (dNHBE) cells were made to order by MatTek Corporation (Ashland, MA) and arrived in kits with either 12- or 24-well inserts each. dNHBE cells were grown on 6mm mesh disks in transwell inserts. During transportation the tissues were stabilized on a sheet of agarose, which was removed upon receipt. One insert was estimated to consist of approximately 1.2 × 10^6^ cells. Kits of cell inserts (EpiAirway™ AIR-100, AIR-112) originated from a single donor, # 9831, a 23-year old, healthy, non-smoking, Caucasian male. The cells have unique properties in forming layers, the apical side of which is exposed only to air and that creates a mucin layer. Upon arrival, the cell transwell inserts were immediately transferred to individual wells of a 6-well plate according to manufacturer’s instructions, and 1 mL of MatTek’s proprietary culture medium (AIR-100-MM) was added to the basolateral side, whereas the apical side was exposed to a humidified 5% CO_2_ environment. Cells were cultured at 37°C for one day before the start of the experiment. After the 24 h equilibration period, the mucin layer, secreted from the apical side of the cells, was removed by washing with 400 μL pre-warmed 30 mM HEPES buffered saline solution 3X. Culture medium was replenished following the wash steps.

#### Viruses

SARS-CoV-2 strain USA-WA1/2020 was passaged twice in vervet kidney cell derived Vero 76 cells to create the virus stock. Virus was diluted in AIR-100-MM medium before infection, yielding a multiplicity of infection (MOI) of approximately 0.003 CCID_50_ per cell.

#### Experimental design

Each compound treatment (120 μL) and virus (120 μL) was applied to the apical side, and compound treatment only was applied to the basal side (1 mL), for a 2 h incubation. As a virus control, 3 of the cell wells were treated with placebo (cell culture medium only). Following the 2 h infection, the apical medium was removed, and the basal side was replaced with fresh compound or medium. The cells were maintained at the air-liquid interface. On day 5, the medium was removed and discarded from the basal side. Virus released into the apical compartment of the tissues was harvested by the addition of 400 μL of culture medium that was pre-warmed at 37°C. The contents were incubated for 30 min, mixed well, collected, thoroughly vortexed and plated on Vero 76 cells for virus yield reduction (VYR) titration. Triplicate and singlet wells were used for virus control and cell controls, respectively.

#### Determination of virus titers from each treated cell culture

Vero 76 cells were seeded in 96-well plates and grown overnight (37°C) to 90% confluence. Samples containing virus were diluted in 10-fold increments in infection medium and 200 μL of each dilution transferred into respective wells of a 96-well microtiter plate. Four microwells were used for each dilution to determine 50% viral endpoints. After 5 days of incubation, each well was scored positive for virus if any cytopathic effect (CPE) was observed as compared with the uninfected control, and counts were confirmed for endpoint on days 6 and 7. The virus dose that was able to infect 50% of the cell cultures (CCID_50_ per 0.1 mL) was calculated by the Reed-Muench method (14). The day 5 values are reported. Untreated, uninfected cells were used as the cell controls.

## Results

Against MERS, there was an EC_50_ of 3.1 to 3.2 μM for both PT150 and PT156 (EC_90_ not calculated). These were compared to an experimental anti-corona virus compound (M128533) with average EC_50_ of 0.83 μM. M128533 showed little cytotoxicity by visual or neutral red reads, CC_50_ >100 μM. Both PT150 and 156, however, showed significant cytotoxicity to the MA104 cells used for the assay: CC_50_ for PT150 in these cells averaged 3.7 μM and for PT156 averaged 9.2 μM. The reasons for this probably relate, in part, to the significantly different GR structure in this species compared to humans; Vero cells, derived from the same source tissues as MA-104 cells, show no response to dexamethasone (20). This cytotoxicity made confident assessment of the antiviral efficacy of the compounds difficult.

Since the very high MA-104 cell toxicity is not present in human cells, as demonstrated by the PT compounds’ extensive pre-clinical assessments (6, 15–20), the absence of documented tissue toxicities in human clinical trials (6), and, in particular, the absence of toxicity in human epithelial airway cells at a 1 μM dose (3), the subsequent testing of PT150 against SARS-CoV-2 was performed in human bronchial epithelial lining cells. Data for SARS-CoV-2 testing are presented in Table 1. EC_90_ for the control drug, Remdesivir, was 0. 0013 μM. EC_90_ for PT150 in two parallel replicative experiments were 4.25 μM and 6.80 μM, respectively, averaging 5.55 μM. There were no observed cytotoxicities at the highest PT150 dose given of 30 μM or for the highest dose of Remdesivir (1.66 μM).

**Table 1.** Antiviral efficacy: EC_90_ for compound PT-150 (Palisades Therapeutics/Pop Test Oncology LLC) against SARS-CoV-2.

| Test Compounds | Concentration (μM) | <sup>a</sup>Log<sub>10</sub> CCID<sub>50</sub> virus per 0.2 mL | <sup>a</sup>Log<sub>10</sub> CCID<sub>50</sub> virus per 0.2 mL | <sup>b</sup>EC<sub>90</sub> (μM) |
| --- | --- | --- | --- | --- |
| PT-150 | 30 | 3.00 | 3.00 | 5.55 |
|  | 10 | 3.00 | 3.30 |  |
|  | 3 | 4.00 | 4.30 |  |
|  | 1 | 4.67 | 4.50 |  |
| Remdesivir | 1.66 | 2.00 |  | 0.0013 |
|  | 0.17 | 2.50 |  |  |
|  | 0.017 | 3.30 |  |  |
|  | 0.002 | 3.67 |  |  |
| Virus Control |  | 4.30 |  |  |
|  |  | 4.67 |  |  |
|  |  | 5.00 |  |  |

Each well was scored positive for virus if any CPE was observed as compared with the uninfected control. Vero 76 cells were scored on day 5 and confirmed on days 6 & 7.
<sup>a</sup>Titer results from the virus yield reduction assay.
<sup>b</sup>EC<sub>90</sub> = 90% effective concentration (concentration to reduce virus yield by 1 log<sub>10</sub>) determined by regression analysis.

## Discussion

PT150 and PT156 are two compounds in a novel class of hormone receptor modulators with a primary clinical effect of GR blockade of circulating cortisol, but with low level GR agonism and AR antagonism from its direct binding to these receptors (1). Initial studies against MERS coronavirus in cultured MA-104 cells suggested possible antiviral effects, though cytotoxicity in this vervet kidney derived cell line made confident assessment of actual potency difficult. Based on the possibility of antiviral effects against MERS, PT150 was tested against SARS-CoV-2, though in human bronchial epithelial lining cells to avoid the same cytotoxicity. There was a highly significant antiviral response with an average EC_90_ of 5.55 μM (3130 ng/mL).

Data from a previously conducted depression Phase 1 MAD study in humans shows that following a 300 mg oral dose the mean C_max_ concentration was 4100 ng/mL, with a half-life of approximately 10 hours (6). Clinical dosing has been safely obtained up to 900 mg oral dose per day. These data suggest that a therapeutic concentration should be obtainable by daily oral dosing in an acceptably safe regimen.

Since no cytotoxicity was seen at the highest PT150 dose given of 30 μM, additional studies are pending to further evaluate at what levels cytotoxicity might emerge, if any. Likewise, the surprising potency of Remdesivir in this model system requires further exploration. In one other report we have been able to identify, Remdesivir was found to have significantly higher potency in human epithelial airway cells compared to Vero and Calu-3 cells (22). These further studies regarding Remdesivir potency in human cells are also pending.

The antiviral effects of PT150 observed in this study may be related to its capacity for AR antagonism. SARS-CoV-2 infection is mediated in part by cellular expression of TMPRSS2 (6), a molecule which is primarily regulated by AR expression (21, 23). The initiating steps for SARS-CoV-2 depend on priming of its spike protein by TMPRSS2 cleavage, which is therefore considered essential for viral spread and pathogenesis (7). Hoffman et al have recently provided confirmation that a protease inhibitor targeting TMPRSS2 neutralizes viral entry and that SARS-CoV-2 also depends on ACE2 as its receptor for cell entry (6). Modulation of ACE2 expression and function by AR, directly by AR regulation of ACE2 expression itself (24) and/or indirectly by TMPRSS2 cleavage of ACE2, may further augment viral entry (25). Thus, PT150 activity against SARS-CoV-2 may be a result of AR/GR downregulation through inhibition of TMPRSS2 expression and diminished ACE2 activity.

The GR modulating effects of PT150 may also be of clinical benefit. The original clinical trials for depression showed that patients who showed signs of dysregulated cortisol, with elevated serum cortisol levels, most benefited from the drug (6). Recent data indicate the important role that upregulation of cortisol is likely to play in progression of COVID-19 to severe disease (11). The potential importance of cortisol had already been signaled by clinical aspects of patient populations at particular risk for SARS-CoV-2 infection and severe COVID-19 disease; individuals with advancing age (26), obesity and metabolic syndrome (27), and cardiovascular disease (28) also have strong associations for hypercortisolemia (29). Of course, hypercortisolemia leads to immunosuppression, another risk factor for COVID-19 that is confirmed by the increased risk for COVID-19 in patients maintained on immunosuppression for solid organ transplants (30). The viral infection itself may lead to hypercortisolemia and resultant immunosuppression, as was seen for SARS (31). Finally, corticosteroid immunosuppressive therapies for COVID-19 have been associated with poorer outcomes (32). We therefore speculate that in the early stages of infection cortisol-induced, relative immunosuppression in COVID-19 patients could inhibit host protective immune responses and potentiate viral infection. Additionally, PT150 and related compounds have demonstrated potent anti-neuroinflammatory properties (personal communication, Dr. Ronald Tjalkens, Colorado State University) which may further impact on systemic progression and outcomes of COVID-19) (33).

This interactivity with immunomodulatory effects of endogenous cortisol raises both opportunity and caution. We postulate potential additive effects between these different mechanisms to reduce vulnerability to infection and to treat COVID-19. Directly blocking cell entry of virus, PT150 may be prophylactic for viral infection and inhibitory of viral progression once infected since, as previously noted by Hoffman et al, inhibition of TMPRSS2 (inhibiting viral spike protein priming) and ACE2 (the viral attachment receptor) is likely to inhibit viral cell entry (7). Once infected, the indirect systemic immunomodulatory effects of the compound may further inhibit progression and reduce severity of disease, particularly in the early stages of infection and in *non-severe* COVID-19. However, during advanced stage, *severe* COVID-19, characterized by a cytokine storm and other features of hyperimmunity, use of PT150 would likely be contraindicated in that it would interfere with exogenous therapeutic glucocorticoid administration, recently shown to be beneficial in a study of dexamethsone in advanced stage COVID-19 (34, 35).

In view of these considerations, we suggest several potential clinical uses of PT150 in SARS-CoV-2 infection and COVID-19 based on its demonstrated safety profile through extensive Phase 1 and Phase 2 clinical trials for depression and its active IND for use in COVID-19 treatment (6). These older studies involved pulse treatments with PT150 of two weeks on/two weeks off (6). This protocol was utilized to minimize the possibility of Addisonian syndrome from the cortisol blockade or a rebound Cushing syndrome upon withdrawal of the compound. Neither syndrome was seen, confirming the safety of the dosing schedule. Given that the natural course of most SARS-CoV-2 infections is two weeks, this protocol can be readily adapted for treatment of early, non-severe infection, given the roughly two week duration of infection until clearance. It may also be useful for developing prophylaxis regimens for use by essential workers with high risk of viral exposure or other exposed persons

In conclusion, PT150 is a clinic ready, small molecule with a good safety profile and active IND for COVID-19. The relatively low dose needed for achieving corresponding anti-COVID-19 concentrations demonstrated by these in vitro studies of SARS-CoV-2 viral suppression is in keeping with readily obtainable drug concentration levels from previously tested clinical, daily oral doses. Further studies are pending, in particular to expore higher dose concentrations to determine a CC50 for the compound in this experimental model. PT150 has unique, possibly additive effects as a direct anti-viral and as a potential modulator of the immune system. For these reasons, PT150 merits further study as a treatment for non-severe COVID-19 and as a prophylactic for viral infection, alone and in combination with other potential antiviral agents, such as cathepsin-L-inhibitors, as recently suggested by Liu et al (36).

## Acknowledgments

The authors are grateful for guidance and advice provided by Drs. Javier Martinez-Picado (IrsiCaixa AIDS Research Institute), Daniel R. Kuritzkes (Brigham and Women’s Hospital – Harvard Medical School), Dewleen G. Baker (School of Medicine at the University of California, San Diego), Christopher D. Verrico (Baylor College of Medicine), and Jeff Taylor (HIV+Aging Research Project-Palm Springs).

## References

1. Philip Tatman, Anthony Fringuello, Michael Graner, et al. Pan-cancer analysis to identify a novel class of glucocorticoid and androgen receptor antagonists with potent anti-tumor activity. J Clinical Oncology 2020; 38:15_suppl, e15663–e15663

2. Peeters BW, Tonnaer JA, Groen MB, et al. Glucocorticoid receptor antagonists: new tools to investigate disorders characterized by cortisol hypersecretion. Stress. 2004; 7: 233–41.

3. Joshi T, Johnson M, Newton R, Giembycz MA. The long-acting β2 – adrenoceptor agonist, indacaterol, enhances glucocorticoid receptor-mediated transcription in human airway epithelial cells in a gene- and agonist-dependent manner. Br J Pharmacol. 2015; 172: 2634–53.

4. Kesel AJ, Huang Z, Murray MG, et. al. Retinazone inhibits certain blood-borne human viruses including Ebola virus Zaire. Antivir Chem Chemother. 2014; 23: 197–215.

5. Christopher D. Verrico, Marguerite M. Patel, Ashley Xu, Victoria B. Risbrough, Dewleen G. Baker, Thomas R. Kosten. A Phase 1 Clinical Trial to Evaluate Pharmacodynamic Interactions after Oral Coadministration of Alcohol and the Highly Selective Glucocorticoid Receptor Antagonist, PT150. Neuropsychopharmacology 2019; 44: 189–90.

6. Investigator’s Brochure on ORG 34517, Edition No. 12. 2009, unpublished (available under confidential disclosure agreement).

7. Hoffmann M, Kleine-Weber H, Schroeder S, et al. SARS-CoV-2 cell entry depends on ACE2 and TMPRSS2 and is blocked by a clinically proven protease inhibitor. Cell. 2020; 181: 271–80.

8. Ghazizadeh Z, Majd H, Richter M, et al. Androgen regulates SARS-COV-2 receptor levels and is associated with severe COVID-19 symptoms in men. Version 2. bioRxiv. 2020 May 15:2020.05.12.091082. doi: 10.1101/2020.05.12.091082.

9. Bhowmick NA, Oft J, Dorff T, et al. COVID-19 and androgen targeted therapy for prostate cancer patients. Endocr Relat Cancer. 2020 Jun 1:ERC-20-0165.R1. doi: 10.1530/ERC-20-0165.

10. Wambier CG, Goren A. Severe acute respiratory syndrome coronavirus 2 (SARS-CoV-2) infection is likely to be androgen meiated. J Am Acad Dermatol. 2020; 83: 308–309.

11. Tan T, Khoo B, Mills EG, Phylactou M, Patel B, Eng PC, Thurston L, Muzi B, Meeran K, Prevost AT, Comninos AN, Abbara A, Dhillo WS. Association between high serum total cortisol concentrations and mortality from COVID-19. Lancet Diabetes Endocrinol. 2020 Jun 18:S2213–8587(20)30216-3.

12. Heneka MT, Golenbock D, Latz E, Morgan D, Brown R. Immediate and long-term consequences of COVID-19 infections for the development of neurological disease. Alzheimers Res Ther. 2020; 12: 69. doi: 10.1186/s13195-020-00640-

13. Day CW, Baric R, Cai SX, et al. A New Mouse-Adapted Strain of SARS-CoV as a Lethal Model for Evaluating Antiviral Agents in Vitro and in Vivo Virology. 2009; 395: 210–22.

14. Reed LJ, Muench H. A simple method of estimating fifty percent endpoints. The American Journal of Hygiene 1938; 27: 493–497.

15. Bachmann CG, Linthorst ACE, Holsboer F, Reul JMHM. Effect of Chronic Administration of Selective Glucocorticoid Receptor Antagonists on the Rat Hypothalamic-Pituitary-Adrenocortical Axis. Neuropsychopharmacology. 2003; 28: 1056–67.

16. Ang-Bleuel A, Ulbricht S, Chandramohan Y, et al. Psychological Stress Increases Histone H3 Phosphorylation in Adult Dentate Gyrus Granule Neurons: Involvement in a Glucocorticoid Receptor-Dependent Behavioural Response. Eur J Neurosci. 2005; 22: 1691–700.

17. Johnson DA, Grant EJ, Ingram CD, Gartside SE. Glucocorticoid receptor antagonists hasten and augment neurochemical responses to a selective serotonin reuptake inhibitor antidepressant. Biol Psychiatry. 2007; 62: 1228–35.

18. Groenink L, Dirks A, Verdouw PM, et al. CRF1 not glucocorticoid receptors mediate prepulse inhibition deficits in mice overexpressing CRF. Biol Psychiatry. 2008; 63: 360–8.

19. Rice BA, Saunders MA, Jagielo-Miller JE, Prendergast MA, Akins CK. Repeated subcutaneous administration of PT150 has dose-dependent effects on sign tracking in male Japanese quail. Exp Clin Psychopharmacol. 2019; 27: 515–21.

20. Reynolds AR, Saunders MA, Brewton HW, Winchester SR, Elgumati IS, Prendergast MA. Acute oral administration of the novel, competitive and selective glucocorticoid receptor antagonist ORG 34517 reduces the severity of ethanol withdrawal and related hypothalamic-pituitary-adrenal axis activation. Drug Alcohol Depend. 2015; 154:100–4.

21. National Institutes of Health. TMPRSS2 transmembrane serine protease 2 [Homo sapiens (human)] Gene ID: 7113. Updated March 13, 2020. Available at: https://www.ncbi.nlm.nih.gov/gene/7113. Accessed April 2, 2020.

22. Sheahan TP, Sims AC, Zhou S, et al. An orally bioavailable broad-spectrum antiviral inhibits SARS-CoV-2 in human airway epithelial cell cultures and multiple coronaviruses in mice. Science Translational Medicine 29 Apr 2020; 12. DOI: 10.1126/scitranslmed.abb5883.

23. Lucas JM, Heinlein C, Kim T, et al. The androgen-regulated protease TMPRSS2 activates a proteolytic cascade involving components of the tumor microenvironment and promotes prostate cancer metastasis. Cancer Discov. 2014; 4: 1310–25.

24. Chen J, Jiang Q, Xia X, et al. Individual variation of the SARS-CoV-2 receptor ACE2 gene expression and regulation. Aging Cell. 2020 Jun 19. doi: 10.1111/acel.13168.

25. Heurich A, Hofmann-Winkler H, Gierer S, Liepold T, Jahn O, Pohlmann S. TMPRSS2 and ADAM17 cleave ACE2 differentially and only proteolysis by TMPRSS2 augments entry driven by the severe acute respiratory syndrome coronavirus spike protein. J Virol. 2014; 88: 1293–307.

26. Zhou F, Yu T, Du R, et al. Clinical course and risk factors for mortality of adult inpatients with COVID-19 in Wuhan, China: a retrospective cohort study. Lancet. 2020; 395: 1054–1062.

27. Zheng KI, Gao F, Wang XB, et al. Obesity as a risk factor for greater severity of COVID-19 in patients with metabolic associated fatty liver disease. Metabolism. 2020 Apr 19:154244. doi: 10.1016/j.metabol.2020.154244.

28. Mehra MR, Desai SS, Kuy S, Henry TD, Patel AN. Cardiovascular Disease, Drug Therapy, and Mortality in COVID-19. N Engl J Med. 2020 May 1. doi: 10.1056/NEJMoa2007621. [Epub ahead of print]

29. Russell G, Lightman S. The human stress response. Nat Rev Endocrinol. 2019; 15: 525–534.

30. Fernández-Ruiz M, Andrés A, et al. COVID-19 in solid organ transplant recipients: a single-center case series from Spain. Am J Transplant. 2020 Apr 16. doi: 10.1111/ajt.15929.

31. Panesar NS, Lam CW, Chan MH, Wong CK, Sung JJ. Lymphopenia and neutrophilia in SARS are related to the prevailing serum cortisol. Eur J Clin Invest. 2004; 34: 382–4.

32. Russell CD, Millar JE, Baillie JK. Clinical evidence does not support corticosteroid treatment for 2019-nCoV lung injury. Lancet 2020; 395: 473–5.

33. Koralnik IJ, Tyler KL. COVID-19: A Global Threat to the Nervous System. Ann Neurol 2020 Jun 7. doi: 10.1002/ana.25807. Online ahead of print.

34. Horby, P. Etal. Preprint at medRxiv https://doi.org/10.1101/2020.06.22.20137273 (2020).

35. Ledford, H. Coronavirus breakthrough: dexamethasone is first drug shown to save lives. Nature 582, 469 (2020) doi: 10.1038/d41586-020-01824-5.

36. Liu T, Luo S, Libby P, Shi G-P. Cathepsin L-selective inhibitors: a potentially promisng treament for COVID-19 patients.

